# Metabolite-based genome-wide association studies enable the dissection of the genetic bases of bioactive compounds in Chickpea seeds

**DOI:** 10.1101/2025.06.16.659862

**Authors:** Siyue Ruan, Lorenzo Rocchetti, Elena Bitocch, Regina Wendenburg, Chiara Santamarina, Valerio Di Vittori, Tawffiq Istanbuli, Aladdin Hamwieh, Alisdair R. Fernie, Roberto Papa, Saleh Alseekh

## Abstract

Chickpea, is the second most consumed food legume, and significantly contributes to the human diet. Chickpea seeds are rich in a wide range of metabolites including bioactive specialized metabolites influencing nutritional qualities and human health. However, the genetic basis underlying the metabolite-based nutrient quality in chickpea remains poorly understood. Here we dissected the genetic architecture of seed metabolic diversity and explored how domestication shaped the chickpea metabolome. Through UPLC-MS we quantify over 3400 metabolic features in 509 chickpea seeds accessions from three independent multi-location field trials. The metabolite genome-wide association study (mGWAS) detected around 130,000 leading SNPs corresponding to 1890 metabolites across different environments. We further found and functionally validated a gene cluster of three CabHLH transcription factors that regulate soyasaponin biosynthesis in chickpea seeds. Our results reveal new insights on the effects of domestication process on chickpea metabolome. and provide valuable resources for the genetic improvement of the bioactive compounds in chickpea seeds.

## Introduction

Chickpea (*Cicer arietinum*) is one of the most consumed legumes globally, ranking third in total legume production worldwide (FAO, 2022). Moreover, due to the provision of plant-based proteins, and the exploitation of biological nitrogen fixation on agricultural areas, its cultivation represents a valuable solution to safely operate within the planetary boundaries (Richardson et al., 2023). Chickpea seeds are rich in protein, fatty acids, and dietary fiber which contributes significantly to the world’s food security by providing dietary proteins and calories for millions of people. Chickpea is also rich in various bioactive metabolites that have important roles in plant growth, development, and adaptation to environmental stresses (Badhan et al., 2018; Chaturvedi et al., 2024; Gao et al., 2015; Kaur and Prasad, 2021; Kudapa et al., 2024; Kumar et al., 2019; Thakro et al., 2023). In addition, many metabolites such as vitamins, flavonoids, phytosterols, and carotenoids provide essential resources for human health as foods, nutrients, and medicines (Gupta et al., 2017; Kaur et al., 2019; Kaur and Prasad, 2021; Sedlakova et al., 2023). Over the past decades, the integration of metabolic profiling and other omics tools provides an effective approach for identification of gene function and elucidation of metabolic pathways (Alseekh et al., 2021b; Shen et al., 2023). Exploring metabolite diversity and its underlying genetic variation provided new insights into crop domestication and breeding as well as germplasm conservation (Alseekh et al., 2021b; Fernie, 2021; Rho et al., 2019). In chickpea, despite the recent progress has been made at the genomic level and characterization of key yield associated traits (Thakro et al., 2024; Varma Penmetsa et al., 2016) the genetic bases underlying the metabolic variation and how the seed metabolome was shaped by domestications are largely unknown. It has been shown that domestication and subsequent diversification represent adaptations to human-built environments and evolutionary forces that shape general phenotypic diversity in chickpea (Varma Penmetsa et al., 2016; Varshney et al., 2021; Varshney et al., 2019). During crop domestication and expansion, metabolites which facilitate cultivation and improve the appearance and taste of food grains were intentionally selected for by farmers (Alseekh et al., 2021a; Bellucci et al., 2021). In some cases, however, the metabolite composition may have been selected due to the closed proximity of the genes or quantitative trait loci (QTL) for specialize metabolite biosynthesis to those regulating traits as nutrients (Alseekh et al., 2015; Ferrao et al., 2020). Secondary metabolites serve key roles in defense and signaling in plants. They contribute to adaptive traits and ecological fitness, including defense mechanisms, tolerance to biotic and abiotic stresses (Chaturvedi et al., 2024; Kudapa et al., 2024; Kumar et al., 2019; Peng et al., 2017; Templer et al., 2017). Plant secondary metabolites are also rich dietary sources of nutrition, energy and medicine. For example, as one of the main bioactive components in legumes, flavonoids served as signaling molecules for rhizobia interactions (Leoni et al., 2021). They additionally have beneficial effects for human health due to their antioxidant activities, and reduce the risk of cancer, heart and cardiovascular diseases (Guardado-Felix et al., 2017; Segev et al., 2010; Wang et al., 2008). The domestication of chickpea led to significant alterations in the composition and abundance of secondary metabolites, which in turn influence both nutritional value and sensory properties (Thakro et al., 2023; Thakro et al., 2024). Understanding the metabolic shifts during domestication is therefore essential for optimizing dietary quality and enhancing nutrient density.

Genome-wide association study (GWAS) and metabolic QTL (mQTL) mapping have been successfully applied to both model plants and major crops to understand the metabolic natural variations and characterize complex quantitative traits (Fang and Luo, 2019). Moreover, significant effort has been expended in order to associate metabolite compositions with quality traits and responses to various environmental stress (Chen et al., 2018; Chen et al., 2020b; Ferrao et al., 2020; Peng et al., 2017; Templer et al., 2017). In this manner, various biosynthetic and regulatory genes of certain metabolites have been cloned in major crop plants, such as flavonoids in rice (Peng et al., 2017), steroidal alkaloids in tomato (Alseekh et al., 2015; Alseekh et al., 2017; Zhu et al., 2022; Zhu et al., 2018) and saponins in soybean (Berhow et al., 2002; Chung et al., 2020; Sundaramoorthy et al., 2019; Tsuno et al., 2018). In Chickpea, despite the recent progress has been mad at the genomics levels and characterization of important phenotypic observations, the genetic bases underlying the metabolic variation have not preformed yet. However, the limited exploration of these metabolomic changes in chickpea has constrained breeding efforts aimed at enhancing the content of bioactive and nutritionally important compounds.

In this study, we used liquid chromatography–mass spectroscopy (LC–MS) to perform comprehensive metabolic profiling of seeds from 509 chickpea lines cultivated across multiple environments. By utilizing metabolomic GWAS (mGWAS), we dissected the genetic bases of metabolic diversity in chickpea seeds. Our research constitutes a valuable resource for understanding chickpea metabolome and reveals new insights into genetic and molecular architecture of metabolic biosynthesis that can enable crop genetic improvement. We described the metabolic variations across *desi* and *kabuli* chickpea market types and showed how the chickpea seed metabolome was altered by selection during adaptation to different environments after domestication. By employing a combined functional genomics and metabolomics, we deciphered the genetic architecture of metabolite quantitative traits in chickpea. Numerous loci and candidate genes for several compound classes were identified; in particular, we found that a gene cluster of three bHLH transcription factors, regulates soyasaponin metabolism in chickpea seeds. The metabolomics diversity and mGWAS results constructed in this study provide valuable data resources for further studying the biosynthetic pathways and regulatory mechanisms of important natural products in chickpea.

## Results

### Specialized metabolite profiling of 509 Chickpea accessions

In previous research we assembled a collection of 531 chickpea accessions and preformed high-resolution genotyping by whole-genome sequencing (WGS) (Rocchetti et al., 2024). Using 701,899 chromosome-wide single nucleotide polymorphisms (SNPs), we assessed the genetic diversity and population structure and showed that there are no a clearly defined number of ancestral populations, suggesting the presence of continuous genetic variation or extensive admixture among the accessions (Fig.1ab). Neighbor-joining (NJ) tree analysis grouped the accessions into three major clusters, reflecting a separation among the two chickpea types (Fig.1a). Among these groups, several chickpea accessions originating from Europe and the Americas exhibited recent divergence. Notably, almost all of these accessions belong to the *kabuli* type, highlighting significant effects of human selection and cultivation practices on the domestication and geographic distribution of chickpea (Fig.1a).

**Figure 1.**
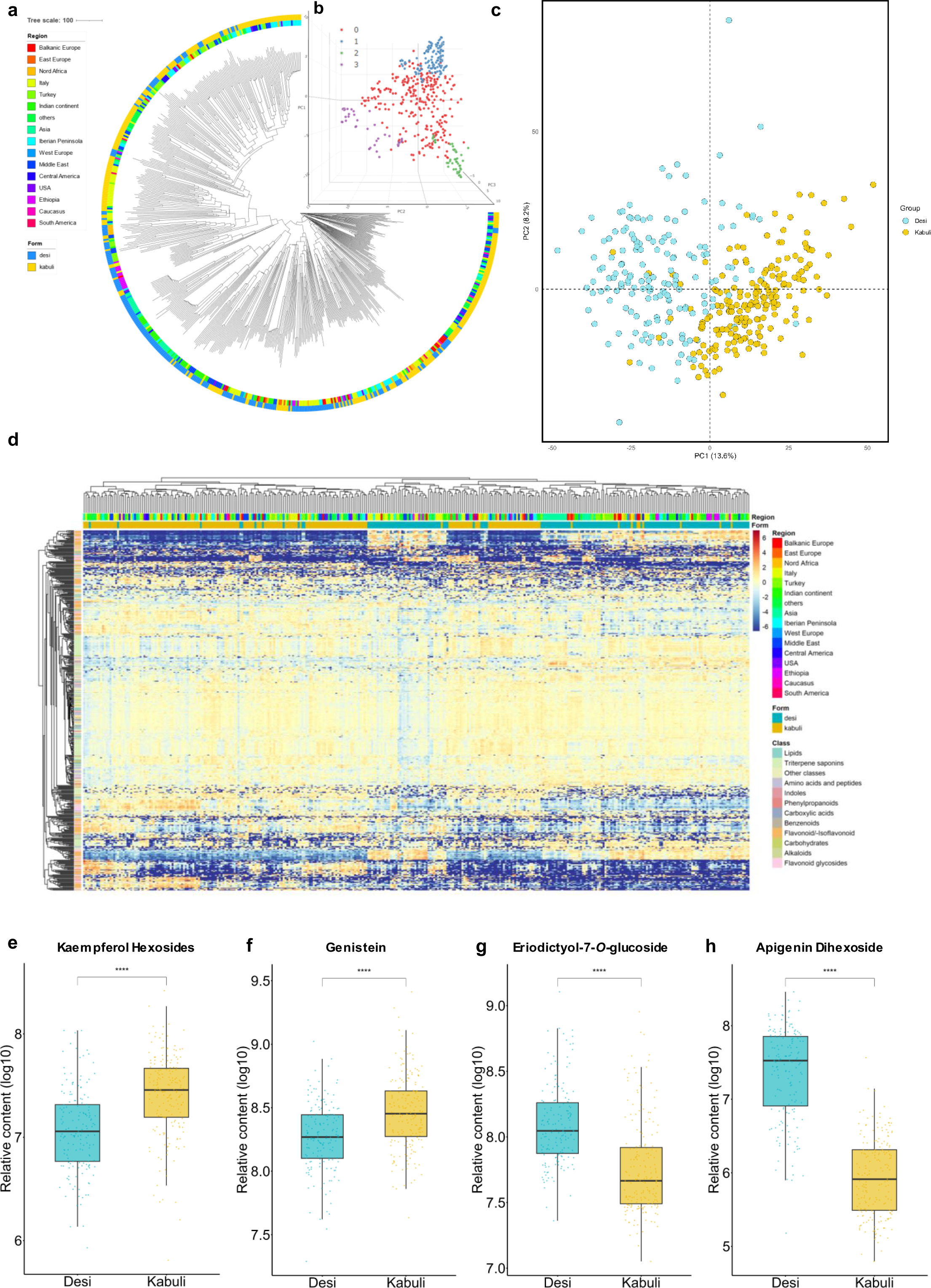
Comprehensive analysis of genomic and metabolic diversity of 509 chickpea accessions. (a) Neighbor-joining (NJ) tree analysis of 531 chickpea accessions based on 700 thousand SNPs. The outer circle represents the different forms of chickpea seeds, and the inner circle represents the origin of chickpea accessions (b) 3D Principal component analysis (PCA) of 531 chickpea accessions using 700 thousand SNPs. 0, 1, 2, 3 represent population structure groups (K=3). (c) Principal component analysis (PCA) of metabolite diversity of chickpea seeds. Score plot with the highest two PCs describing 13.6% and 8.2% of the variation, respectively. Yellow triangle = *kabuli*, light blue triangle =*desi*. (d) Heatmap showing the relative abundancies of 500 annotated metabolites in chickpea seeds. Groups are defined based on the domestication form. (e-h) Relative abundance of highest differential metabolite features across desi and kabuli accessions. The significance of the differences was analyzed by two-tailed t-test. Asterisks indicate level of statistical significance: * p ≤ 0.05.

To explore the chemical diversity of chickpea seeds, around 2,000 seed samples were collected from three independent field trials conducted in Italy (2020 and 2021) and Lebanon (2022). Comprehensive metabolic profiling was preformed using ultra-performance liquid chromatography coupled with tandem mass spectrometry (UPLC-MS/MS) (Alseekh, 2021). After chromatogram processing and filtering, 3461 metabolites were determined across 509 seed accessions. Of which, 771 metabolites were annotated based on common fragmentation and in house metabolite library, while 2610 remained unknown (Fig. 1bc; Table. S1). Results revealed the presence of a wide array of important metabolic subclasses in chickpea seed such as flavonoids. Principle Component Analysis (PCA) was preformed to explore the overall metabolic diversity in the chickpea panel. The first two PCs explain 26.8% of the variation (PC1: 18.8%; PC2: 8%) (Fig. 1b), in addition heatmaps demonstrated the relative changes in seed metabolites across the 509 chickpea accessions (Fig. 1b). Results indicate that the origin of the two major seed form (*desi* and *kabuli)* is the main force driving the metabolomic separation (diversity) across the studied chickpea accessions (Fig.1bc). Further analysis showed that 299 metabolites shared significant differences between *desi* and *kabuli* with more than two-fold relative changes. Among the most abundant and differential metabolite subclass in chickpea seeds was flavonoids, we identified 17 flavonoid derivatives, contains flavonoids, isoflavonoids and flavonoid glycosides. Specifically, most flavonoids were tightly associated in one metabolite cluster, these highly correlated metabolites tended to be of similar structure (e.g., Flavonoid-*3-O*-glycosides and Flavonoid-*7-O*-glycosides) or intermediates of the same biosynthesis pathway, supporting their involvement in similar physiological processes. Such metabolites are known to be associated with the seed coat color and hence are significantly altered between *desi* and *kabuli* types of the chickpea seeds.

To explore the metabolic variation in chickpea shaped by diversification and adaptation following the domestication and spread beyond the area of origin, we selected 290 chickpea accessions located from major cultivation area. Partial Least Squares Discriminant Analysis (PLS-DA) was conducted to elucidate differences in relative metabolite feature intensities among chickpea accessions of distinct geographical origins. (Fig. 2a). Data showed that accessions from Turkey are significant separated from the ones coming from India. Syrian accessions are clustered closer to Turkish accessions in terms of metabolite composition, while African accessions were more similar with the Indian accessions (Fig. 2a). On the other hand, the Central Asian accessions are more dispersed, also being significant separated with the Indian accessions. This observation is consistent with two major chickpea domestication pathways, as the first path illustrates a diffusion to India and East Africa and the second path suggests a diffusion to Central Asia together with the Mediterranean region (Varshney et al., 2021). During the dissemination and adaptation process to different environments, the main altered subclasses of secondary metabolites were flavonoids, epigallocatechin, triterpene saponins and peptides (Fig. 2b-f).

**Figure 2.**
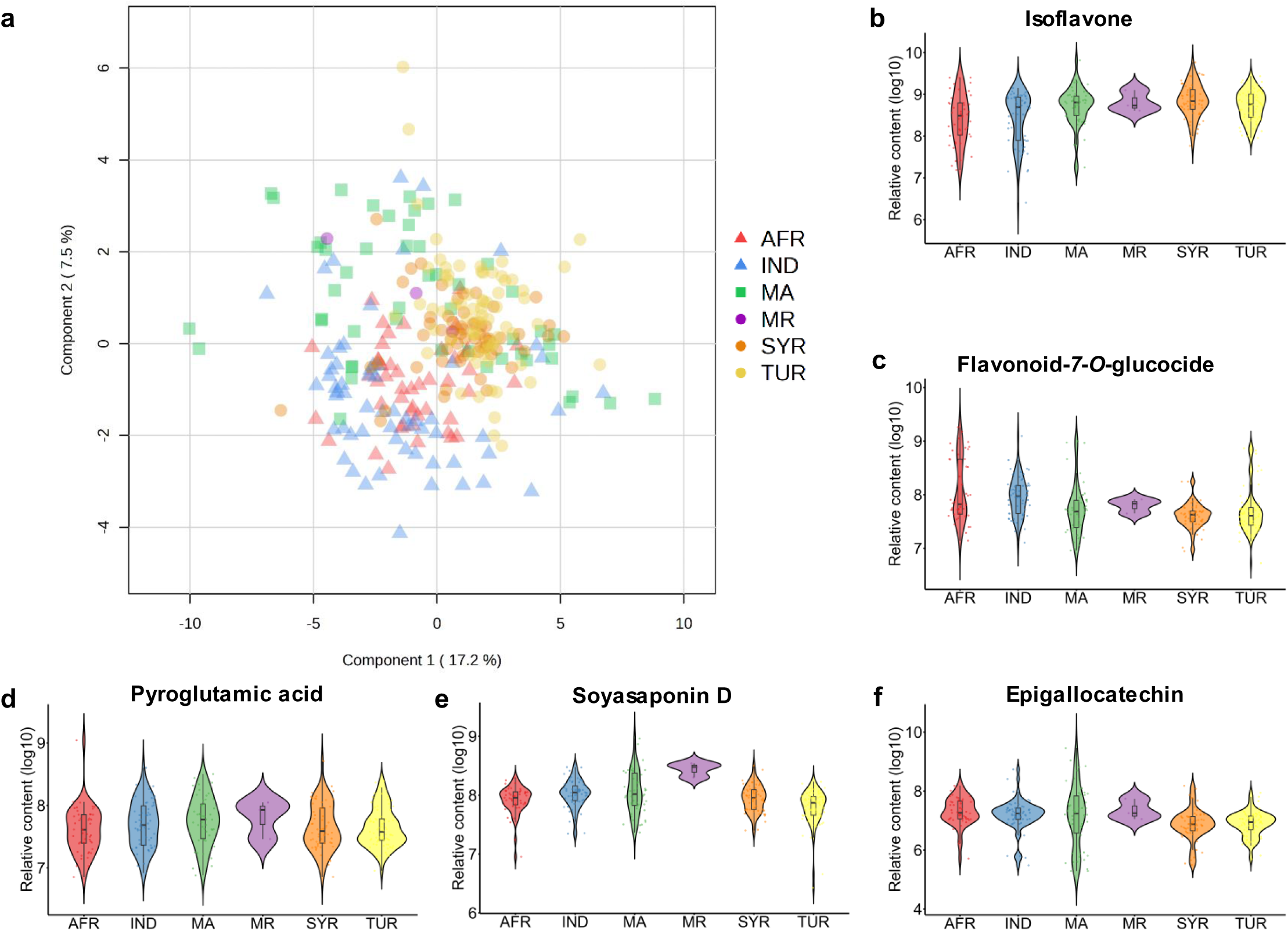
Comprehensive analysis of metabolite-derived domestication distribution of 290 chickpea accessions. (a) Partial Least Squares Discriminant Analysis (PLSDA) of metabolic variation among individuals based on the origin of domestication. (b-f) Effect of domestication on highest differential metabolite compounds in chickpea seeds. Red, AFR, Africa; Orange, SYR, Syria; Yellow, TUR, Turkey; Blue, IND, India; Purple, MR, Middle east Asia; Green, MA, Central Asia.

### Genetic basis of nature metabolic variations in chickpea seed

To identify genomic loci associated with metabolic variations in chickpea seed, we integrated a total of 701,899 high-confidence chromosome-wise SNPs that were obtained from the whole genome sequencing of 509 chickpea accessions (Rocchetti et al., 2024) and subsequently used to perform the mGWAS analysis. We identified 125,054, 133,092 and 127,794 SNPs corresponding to 1,618, 1,700 and 1,890 metabolites in three different experiments, respectively (Table S2). Approximately 46.7%, 49.1% and 54.61% (1890/3461) of the detected metabolites were associated with at least one SNP, with an average of 50.2% per trial. The full list of significant mGWAS association SNP is summarized in Table S2.

The Circos plot displays the number and distribution of the identified significant GWA hits for specialized metabolites in all three environments (Fig. 3). The significant loci were not randomly distributed across the eight chickpea chromosomes, indicating an enrichment of major genes controlling the levels of multiple metabolites in several genome regions. Moreover, the number and distribution of significant associations well reflected the metabolic genetic stability of our population (Fig. 3), indicating a consistent genetic basis of metabolite variation across environments. Based on metabolite subclasses, we classified the annotated metabolites, and replotted the distribution of these SNPs using different metabolic traits (Fig. 3-4). In total, over 40 hotspots were identified, specifically, four major hotspots associated with peptides, oxitripan, flavonoids and soyasaponin derivatives were found on Chr3: 0.69-0.75 Mb, Chr4: 6.3-6.4 Mb, Chr5: 7.26-7.33Mb, and Chr7: 1.22-1.27Mb, with high similarity across the three experiments. In addition, we also identified multiple significant QTL associated with various metabolite classes across different experiments. For instance, we found significant SNPs associated with organic acids aspartate and pyroglutamate in Chr4 and Chr6, respectively (Fig.3-4). Next, candidate genes were identified using genomic windows of 100kb upstream and downstream of SNP markers associated with the metabolic traits. Taking the peptide- associated hotspot on Chr3 as an example, a total of 57 genes were identified within a 60 kb genomic window associated with significant SNP markers. Among these genes, a Ca3g091900 was identified as candidate gene encoding Peptide Transporter 3 (PTR3), a member of the peptide transport family (Karim et al., 2005). Additionally, four other genes in this region were identified as orthologs of probable peptide/nitrate transporters in *Glycine max*, suggesting a potential role in nitrogen or peptide transport mechanisms in chickpea (Fig. 4; Table S2).

**Figure 3.**
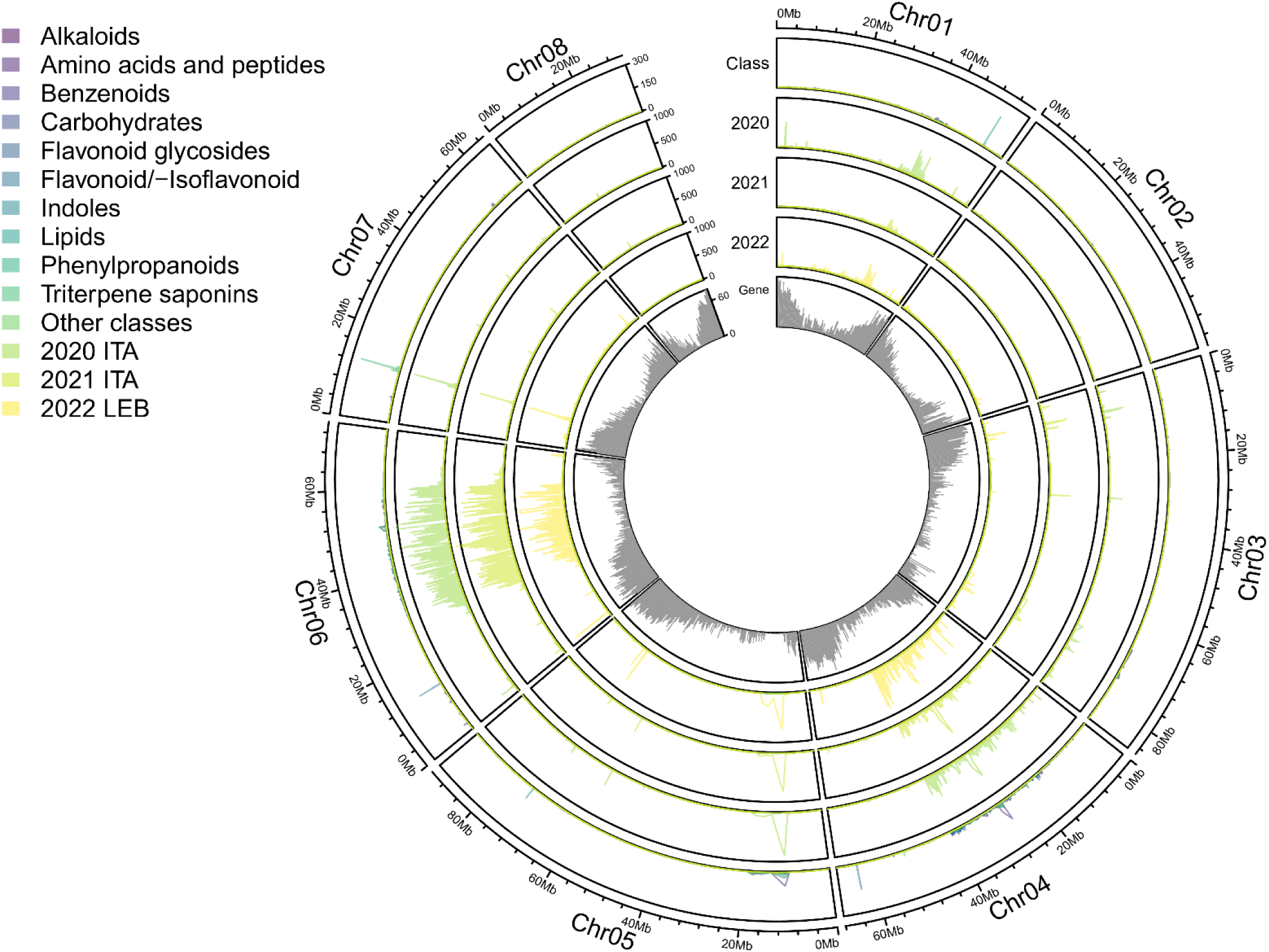
Genome distribution of significant SNPs related to all metabolites in chickpea seed. Chromosomal distribution of the QTL derived from mGWAS represents the combined results from three independent seasons (2020, 2021 and 2022). The circles from outside to center represent different metabolite subclasses and seasons (2020 in Italy, 2021 in Italy and 2022 in Lebanon). The peaks indicate the threshold of hotspot, represented as the number of mGWAS signals within an interval of single gene. Colors indicate different metabolite subclasses. The inner circle specifies the gene density of chickpea genome. All significant SNPs associated with the metabolic traits are listed in Table S2.

**Figure 4.**
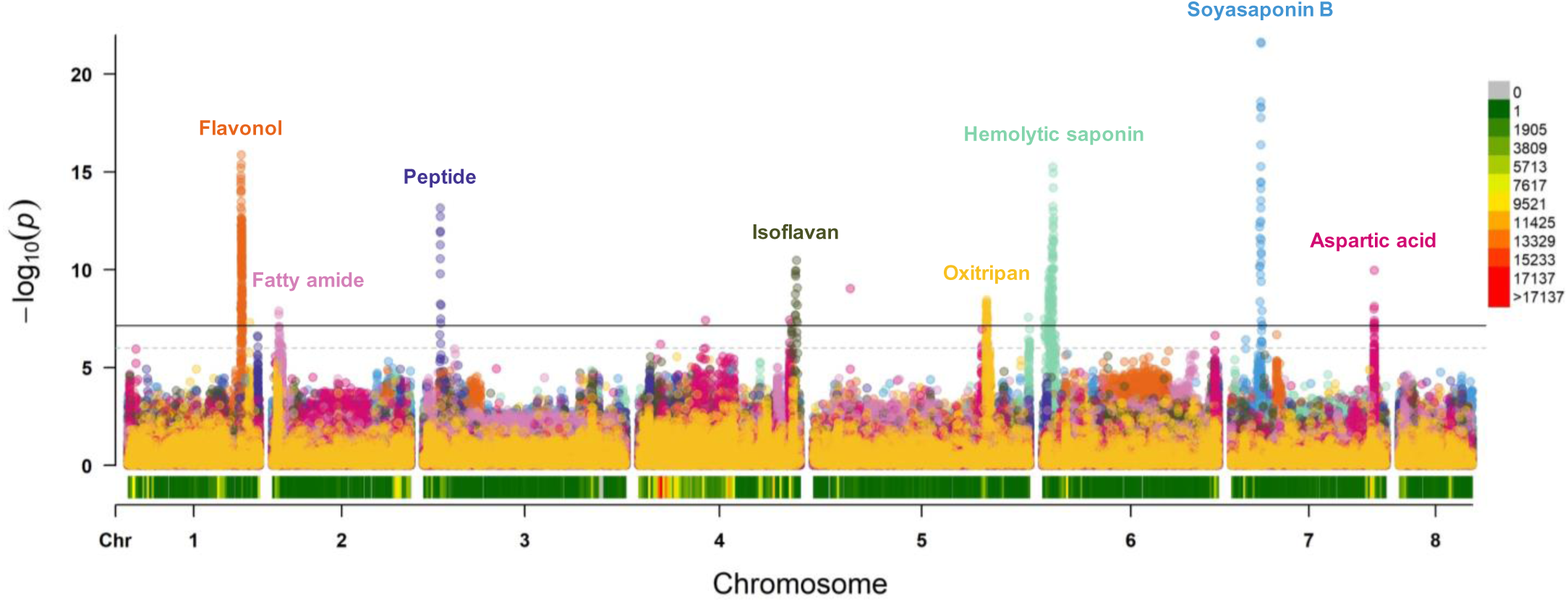
Genome-wide detection of selected metabolite subclasses in chickpea seed. Manhattan plots for soyasaponinB, oxitripan, peptide, flavonol, isoflavane aspartic acid, fatty amide and hemolytic saponin. Complete GWAS results are represented in Table. S2.

To evaluate if there are any significant associations specific to the two forms of chickpea seeds, we next conducted mGWAS analysis independently on *kabuli* and *desi* chickpea. Results indicate that, 829, 807 detected metabolites were significantly associated with QTL in *desi*, and *kabuli* seeds, respectively. For instance, the major QTL associated with soyasaponin derivatives in chr.7 consistently demonstrated high significance in both seed types. In *desi*, SNPs associated with peptides and flavonoids exhibited a higher level of significance. All significant GWA hits for specialized metabolites in the two chickpea types were organized and represented in a Circos plot (Fig. S1; Table S3).

Furthermore, to determine whether the above observations are specific to metabolic traits or broadly applicable to other traits, we conducted GWAS analysis on 28 phenotypic traits. Six traits showed significant GWA hits in a continuous genome region on chromosome 4, suggesting that the observed genetic associations to some of the phenotypic traits are likely driven by underlying genetic differences between seed type varieties (Fig. S2; Table S4).

### Functional Interpretation of mGWAS results for oxitripan

We identified genomic region on chromosome 5 strongly associated with oxitriptan (5- hydroxytryptophan) and indole-based compounds, the 700-kb genomic region harbored 108 significant SNPs. Within the linkage region, the highest SNP was located in Ca5g209200 gene annotated as trehalose-phosphate phosphatase J-like gene (Fig.5ab). The leading SNP for oxitriptan exhibited a modest effect size, seed content of oxitriptan was significantly higher in the accessions harboring the A-allele (Fig.5c). By analyzing the functional annotations of all candidate genes in this genomic region, we identified a MYB TF Ca5g210400, homologous to MYB64 in *A. thaliana*, which is shown to be essential for the regulation of female gametophyte development (Rabiger and Drews, 2013), suggesting that this MYB TF may similarly regulate seed development in chickpea. Furthermore, we identified Ca5g220500 gene within 500 kb of the significant SNPs annotated as tryptophan aminotransferase-related protein, which directly play a role in oxitripan biosynthesis (Fig. 5f). Tryptophan is an essential amino acid that plays a crucial role in various biological functions both in plant and animal (Kaluzna-Czaplinska et al., 2019), its derivatives, such as oxitriptan, serotonin, and tryptamine also have specialized functions including (Erland et al., 2016; Lemaire and Adosraku, 2002; Negri et al., 2021; Sun et al., 2025). The metabolic profiling on chickpea seeds collected at different development stages (see materials and methods) showed that the abundance of these metabolites is variable across the seed development in chickpea with significant positive correlation between tryptophan and oxitriptan content, indicating a metabolic linkage in this pathway (Fig. 5e-f).

**Figure 5.**
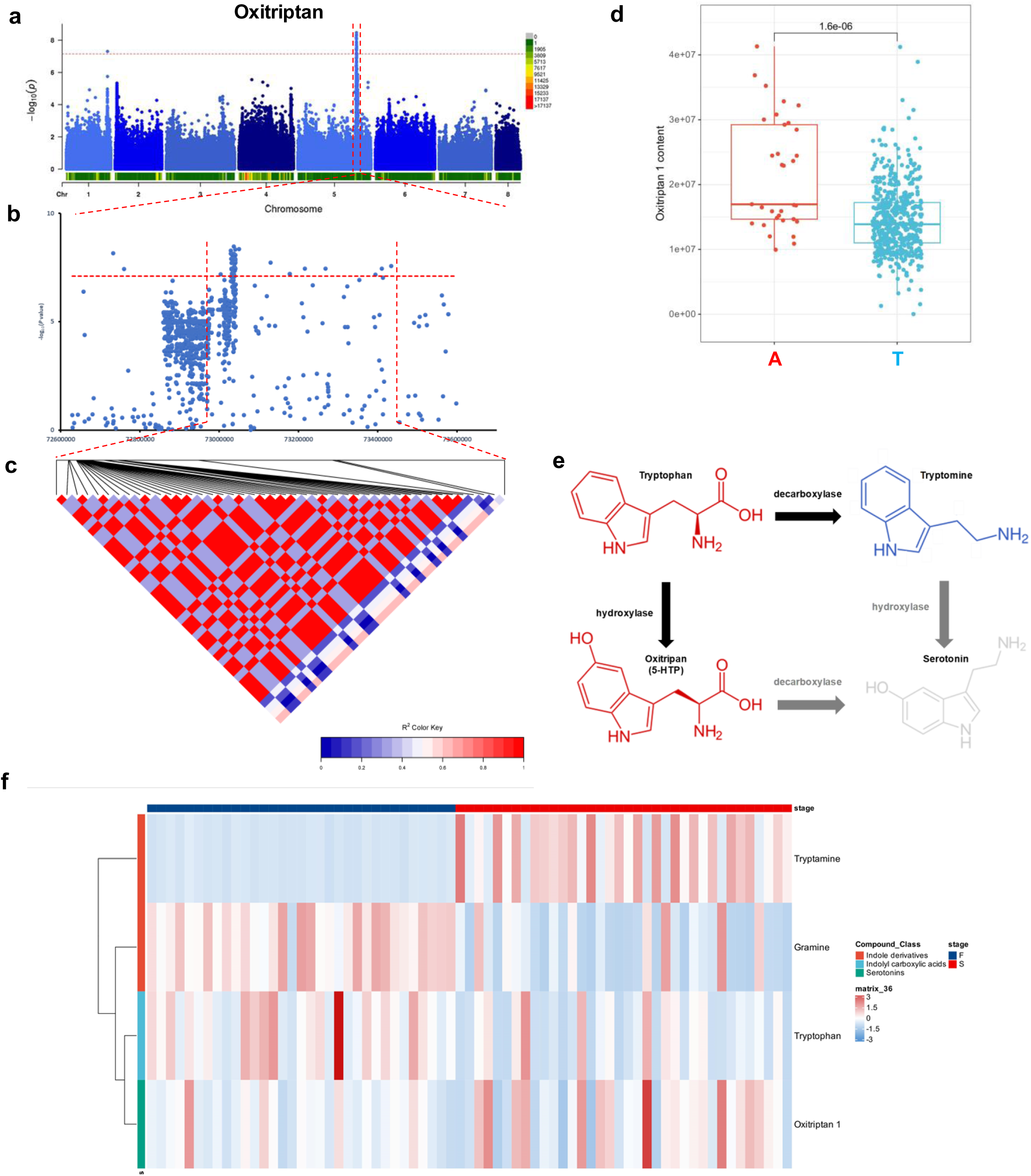
Functional annotation of genes associated with oxitripan variation in chickpea seeds. (ab) Manhattan plot displaying the mGWAS results of Oxitriptan. (c) LD heatmap surrounding the associated peak for Oxitriptan on chromosome 5. (d) Boxplot displaying the difference in relative contents of Oxitriptan between different lines carrying different alleles based on the single likely causative SNP 5:72732870. (e) Putative biosynthetic pathway of serotonins in chickpea. Colors show the correlation among the compounds. Red, positive correlation; Blue, negative correlation. (f) Heatmap of representative indole-based compounds in chickpea seeds during different growth periods. F, Flowering stage; S, Senescence stage. The significance of the differences was analyzed by two-tailed *t*-test.

### Functional characterization of CabHLH TFs that regulate soyasaponins in chickpea seeds

Soyasaponins are oleanane-type triterpenoid saponins derived from β-amyrin, the biosynthesis pathways in legumes begin with cyclisation of the common precursor 2,3- oxidosqualene by oxidosqualene cyclases (OSCs) into various triterpene scaffolds (Biazzi et al., 2015; Chung et al., 2020; Li et al., 2021).

In this study we identified a hotspot mGWAS at chromosome 7 that influenced a wide range of soyasaponin derivatives in chickpea seeds across all three experiments (Fig. 3; Table S2). To identify candidate genes responsible for this locus, we deeply investigated some of the more strongly influenced saponins derivatives e.g., Soyasapogenol B-DDMP- glucuronic acid-hexose-dehexose (DDMP-glcA-hex-dehex) and other soyasaponin B group derivatives. For instance, the leading SNP for DDMP-glcA-hex-dehex (SNPs (- log10(p) = approx. 21)), showed the largest effect size of the leading SNP (Fig. 6ab). Our data indicated that the soyasaponin B group derivatives were more altered with higher soyasaponins content in the accessions containing the T-allele (Fig. 6d). Examining the genes within 500 kb of the significant SNPs lead to identification of a cluster of three bHLH TFs (Table S2). To check the similarity of these TFs to other related plant species, we used full-length protein sequences of the homologous bHLH TFs from four different legume species to construct a phylogenetic Tree (Fig. 6e). The phylogenetic analysis revealed high similarity of the identified bHLH TF cluster across all four species, interestingly, the cluster (bHLH143300) contains one of the *TSAR* (TRITERPENE SAPONIN BIOSYNTHESIS ACTIVATING REGULATOR) genes, which were shown to boost triterpene saponin biosynthesis, and activate genes of the precursor mevalonate pathway in *M. truncatula* (Fig. 6e).

**Figure 6.**
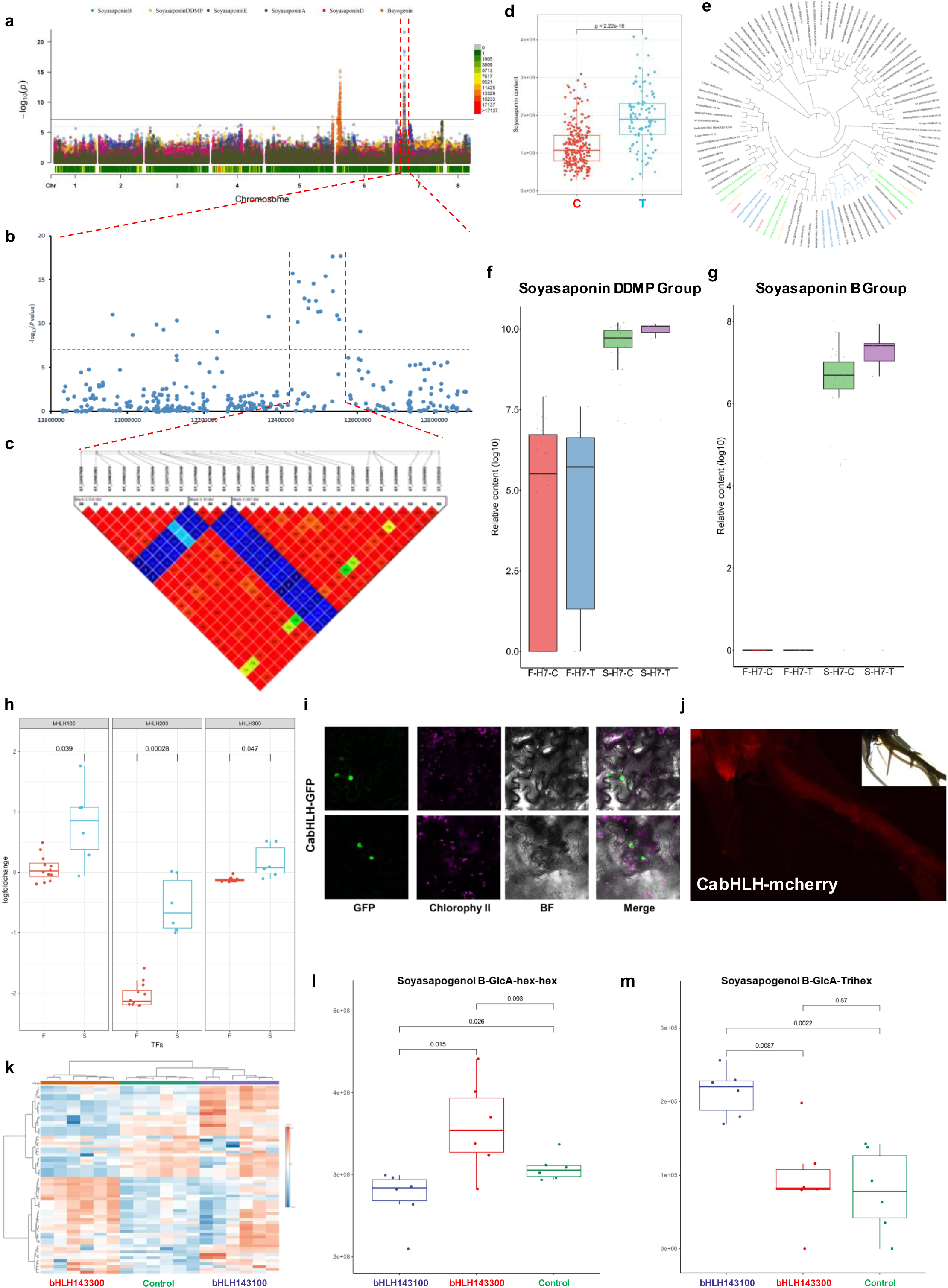
Functional annotation of genes responsible for variation in Soyasaponin content. (ab) Manhattan plot displaying the mGWAS results of annotated saponins in chickpea. (c) LD heatmap surrounding the associated peak for soyasaponins on chromosome 7. (d) Boxplot displaying the difference in Soyasapogenol B-glcA-hex-dehex content between different alleles based on the single likely causative SNP 7:12547295. (e) Phylogenetic tree analysis of CabHLH and bHLH TFs based on the ortholog in four legume species. The approximately maximum- likelihood tree was built by using seeded guide trees and HMM profile-profile techniques to generate alignments between whole length protein sequences. Red, 3 CabHLH TFs; Green, homologous in *Glycine max*; Blue, homologous in *M. truncatula*. (fg) Representative soyasaponin B in chickpea seeds during different growth periods. H7 means the accessions selected based on the harplotype of QTL7. F, Flowering stage; S, Senescence stage. (h) Relative expression of 3 CabHLH TFs in chickpea seed during different growth periods F, Flowering stage; S, Senescence stage. j, Subcellular localization of overexpression of 2 CabHLH TFs in transient tobacco leaves. Confocal images of CabHLH TFs with C-terminal GFP co-expressed in transient tobacco leaves. GFP signals of bHLH-GFP fusion proteins localized in cell nucleus. GFP, fluorescence of bHLH-GFP or eGFP; Chlorophy II, chloroplast autofluorescence; Merged, merged image of GFP and chloroplast; BF, bright field. (j) Observation of mcherry fluorescence in transgenetic hairy roots with CabHLH. (k) Heatmap showing the relative abundancies of 30 soyasaponins in overexpression lines. Groups are defined based on different overexpression lines. (lm) Increased levels of soyasaponin B derivatives in CabHLH overexpression chickpea plants compared with the control (empty vector) in hairy root. The significance of the differences was analyzed by two-tailed *t*-test.

To further explore the genetic basis of soyasaponins in chickpea and further support the mGWAS results, we selected 20 chickpea accessions and preformed transcriptomic and metabolomic profiling across different chickpea tissues at three developmental stages (Fig. S3; Table S5). Results showed that soyaponins are mainly accumulated and distributed in the root and seed, with very low abundance in other tissues. The accumulation of soyasaponins in chickpeas gradually increases across plant development in most tissues, with levels varying depending on the tissue type and specific soyasaponin group. For instance, the data showed that soyasaponins accumulate in chickpea root at very early development stage, and further increase in later growth periods and eventually remain at a higher level (Fig. S4). Furthermore, the life- cycle experiment, demonstrated that the expression of CabHLH TFs was significantly correlated with metabolite levels, specifically the accumulation of certain soyasaponins in chickpea seeds across different growth stages (Fig. 6f-i) supporting the hypothesis that these TFs regulate soyasaponins content in chickpea.

In order to validate the function of CabHLH transcription factors (TFs) in soyasaponin biosynthesis, transient overexpression assays were performed in *N. benthamiana*. Subcellular localization was assessed using a GFP-tagged construct driven by the cauliflower mosaic virus (CaMV) 35S promoter. Fluorescence microscopy revealed that GFP signals were predominantly localized in the nucleus, indicating nuclear localization of the TFs (Fig. 6j). Metabolic profiling of transformed tobacco leaves revealed significant alterations in a distinct cluster of metabolites, characterized by later elution time (RT> 7 min) and mass-to-charge ratios (*m/z*) > 300. Multivariate statistical analysis demonstrated a clear metabolic divergence between the overexpression lines and empty vector controls in transient expression assays (Fig. S5; Table S6).

To further validate this finding, we introduced overexpression constructs (A. rhizogenes K599) for the transcription factors (TFs) into hairy roots of both Phaseolus vulgaris and Cicer arietinum (Fig. 6n; Fig. S6; Table S7), followed by quantification of soyasaponin content in the transgenic roots. UPLC-MS metabolic profiling of the transformed hairy roots showed that 30 annotated soyasaponins exhibited the most significant variation between the CabHLH hairy roots overexpression lines and empty-vector lines (Fig. 6k). Among these, compounds that demonstrated significant accumulation were predominantly characterized by the presence of multiple sugar decorations, such as Soyasapogenol B-glcA-hex-hex-hex/dehex, which closely mirrors the results obtained from the tobacco leaf overexpression experiment.

Furthermore, to reconstruct the biosynthetic pathway of triterpene saponins in chickpea (Fig.7), we performed BLAST searches using the full-length protein sequences of key genes involved in soyasaponin biosynthesis, such as cytochrome P450 monooxygenase (P450s) and UDP-dependent glycosyltransferases (UGTs) from *M. truncatula* and *G. max*, to identify the orthologous genes in chickpea. During soyasaponin biosynthesis, the glycosylation of aglycones is predominantly catalyzed by UGTs, which play a crucial role in determining the structural diversity and bioactivity of soyasaponins. Here, we speculate that the CabHLH TFs affect the biosynthesis of macromolecular soyasaponins by regulating specific UGTs (Fig. 7).

**Figure 7.**
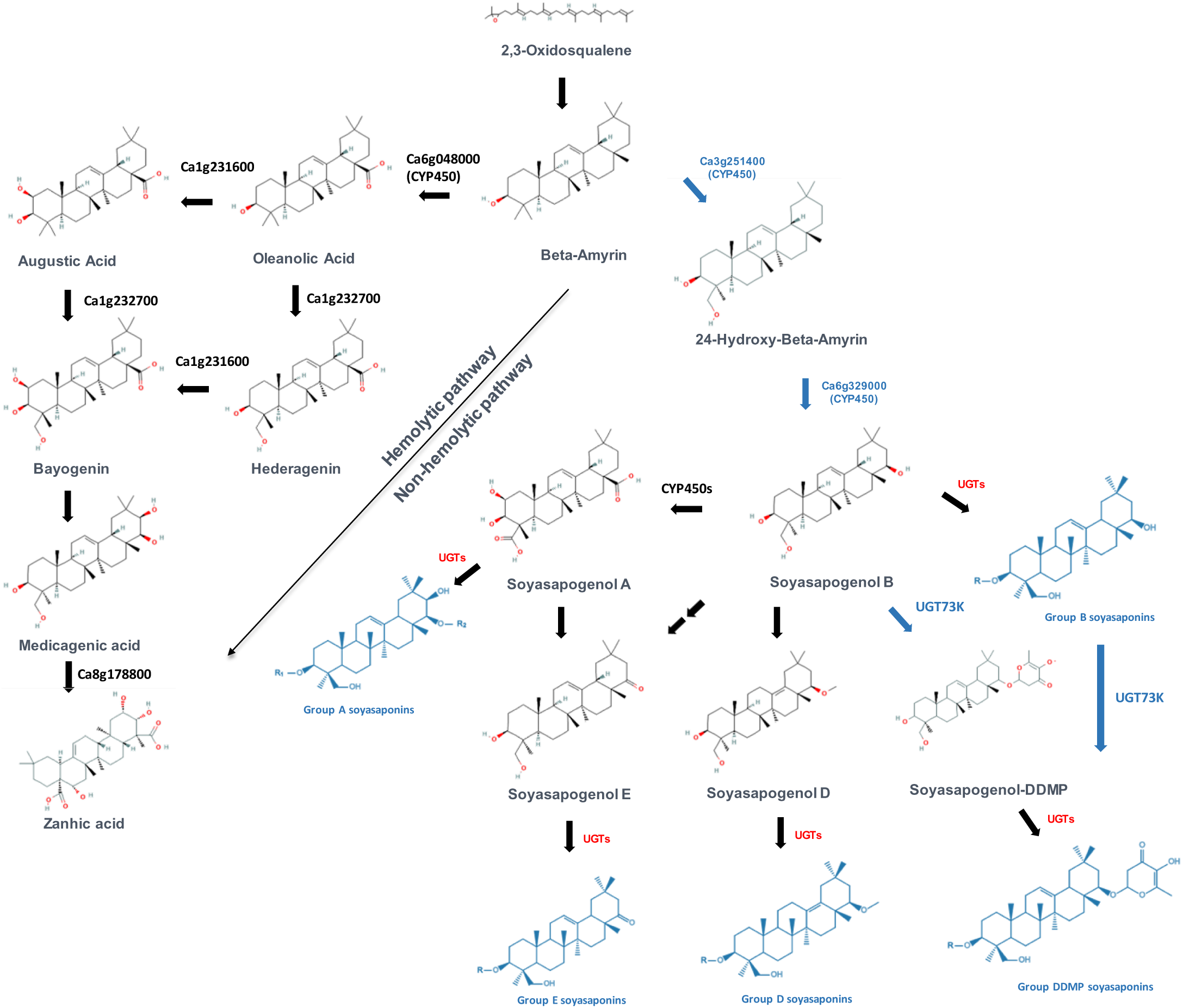
Putative biosynthetic pathway of triterpene saponins in chickpea. Zanhic acid and soyasaponins are oleanane-type triterpenoid saponins, derived from β-amyrin. Genes in red are predicted based on homologous on *M. truncatula* and *G. max*. Saponin compounds and genes inblue are detected and annotated in this paper. Genes in red are predicted to be homologous of UGTs.

## Discussion

Chickpea is one of the most consumed leguminous crops and contributes substantially to the human diets (Gupta et al., 2017; Kaur et al., 2019; Kaur and Prasad, 2021). Seed quality is significantly influenced by the accumulation of various storage bioactive compounds including specialized metabolites (Bulut et al., 2023; Dominguez-Arispuro et al., 2018). Therefore, determination the genetic bases of metabolites content is a fundamental step to develop high- nutritional quality varieties to meet human needs through integration of multi-omics with traditional breeding.

GWAS is an efficient and widely used to identify genetic loci controlling agronomic and metabolic traits (Atwell et al., 2010; Buckler et al., 2009). In recent years, the application of metabolic GWAS studies has focused on dissecting the genetic architecture underlying the biosynthesis regulation of metabolic pathways of many crop species (Alseekh et al., 2021a; Chen et al., 2020a; Ding et al., 2023; Fang and Luo, 2019; Lin et al., 2023; Lu et al., 2021; Zhao et al., 2024). However, the application of this approach to investigate the seed metabolic diversity in chickpea has not yet been previously performed. In the present study, we quantified over 3400 metabolite features in seeds of 509 chickpea accessions and identified 8 major significant associations via metabolic genome-wide association mapping. To demonstrate our finding, we identified and validated a cluster of bHLH transcription factors that regulate the soyasaponin content in the chickpea seed.

The human-guided domestication and subsequent worldwide spread of food crops altered a wide range of agronomic and metabolic traits (Alseekh et al., 2021a; Alseekh et al., 2021b). In chickpea for instance the preference of light-colored (*Kabuli*) and dark-colored (*desi*) is one of the key target traits selected during domestication and artificial selection. Indeed, deciphering the underlying genetic and molecular mechanisms governing the seed types variation have been well studied. For example, QTL mapping was used to identify the genetic basis underlying variations in seed color and seed coat morphology (Thakro et al., 2024). In this study, we identified considerable metabolic diversity between the two chickpea types, with significant variations in the content of flavonoid derivatives and peptides exhibiting bioactivity. The flavonoid metabolic pathway, which modulates seed color, is a potential target of human-driven domestication and selection. Furthermore, Flavonoid-dominated metabolite profiles exhibited highly conserved metabolic diversity, suggesting the existence of coordinated regulatory frameworks influencing spatial and genotypic distribution. Notably, chickpea varieties from India and the Turkey–Syria region exhibit significant metabolic separation, which is consistent with the two main routes of geographic differentiation observed in chickpea domestication (Varshney et al., 2021; Varshney et al., 2019). These findings provided valuable insight into how chickpea adapts to diverse ecological environments by modulating the biosynthesis of bioactive metabolites. However, the impact of these processes on seed nutritional quality and metabolomic profiles remains largely unaddressed.

As a major source of saponin in human diet, chickpea seeds contain a large amount of saponins. Saponins play an important role in plant defensive system as bioactive barriers, but also determine plant nutritional and medicinal qualities (Serventi et al., 2013; Sharma et al., 2023; Zhu et al., 2016). In legume species, soyasaponins influence microbial communities in the rhizosphere. Some studies suggest soyasaponins modulate root– microbe interactions by altering root exudate composition (Korenblum et al., 2020; Leoni et al., 2021; Tsuno et al., 2018). The root of legume releases soyasaponins into the soil can alter microbial community structure and influence nutrient cycling and allelopathic interactions (Tsuno et al., 2018). Therefore, characterizing the regulatory networks moderating saponin content will advance the functional characterization of the biosynthetic pathway in chickpea and facilitate its utilization. The transcriptional regulation of saponin biosynthesis is a multilayered process involving a dynamic interplay of TFs, which have been identified in several plant species. For example, the TRITERPENE SAPONIN BIOSYNTHESIS ACTIVATING REGULATOR (TSAR) TFs regulate saponin biosynthesis in Medicago (Mertens et al., 2016). Moreover, reports indicate that in some medicinal plant species, saponins biosynthesis pathway is also regulated by MYB TFs and ERF TFs (REF). The comprehensive characterization of bHLH TFs in legumes highlights their integral roles in transcriptional regulation associated with development, environmental stress adaptation, and secondary metabolism (Suzuki et al., 2021). The differential expression patterns of bHLH TFs in response to abiotic stress and during symbiotic interactions suggest their involvement in regulatory networks that modulate key physiological responses. Importantly, several bHLH TF members are putatively implicated in the transcriptional control of soyasaponin biosynthesis, while ortholog genes distributes similarly in different legume species. In this study, the functional role of bHLH TFs in regulating soyasaponin biosynthesis was validated in chickpea. Hairy root transformation in chickpea and common bean, along with transient overexpression assays in *N. benthamiana*, demonstrated that a group of CabHLH TFs involved in the soyasaponin biosynthesis pathway in chickpea. Deciphering the tissue-specific expression and functional dynamics of these CabHLH TFs will advance our understanding of the molecular mechanisms encouraging soyasaponin accumulation. Expression profiling indicated that the expression of CabHLH TFs and UGT73like & UGT91like were highly consistent across multiple tissues, while the correlation is supported by the hairy root experiments, where transgenic lines showed higher content of soyasaponins with multiple sugar chains. Though non-nutritive in the traditional sense, soyasaponins contribute significantly to human health through their cholesterol-lowering, anticancer, antioxidant, anti-inflammatory, and immunomodulatory properties (Zhu et al., 2016). Furthermore, such insights have significant implications for crop improvement strategies aimed at enhancing stress resilience and optimizing the nutraceutical quality of chickpea through molecular breeding or biotechnological interventions.

In summary, this study establishes a comprehensive foundation for integrative metabolomic and population genomic analyses in chickpea, with implications for crop breeding. Through untargeted metabolite profiling, we identified more than 3400 kinds of metabolite features and developed a metabolome database encompassing 771 annotated metabolic compounds. This resource enables high-resolution GWAS, facilitating the dissection of metabolic population structure and the identification of genes associated with metabolic variation. Furthermore, we systematically analyzed the geographical and morphotype-specific metabolic differences in chickpea seeds, revealing potential associations between metabolic divergence and germplasm differentiation along migration routes. These findings will support breeders in effectively utilizing genetic diversity and candidate genes linked to bioactive compounds, accelerating the development of nutritionally chickpea cultivars and contributing to improved agricultural productivity and sustainability.

## Materials and Methods

### Plant Material of three field experiments

The chickpea GWAS panel used in this study consists of 509 diverse lines selected from the chickpea collection obtained by single-seed descent and developed within the EMCAP and INCREASE projects with the aim of representing the global diversity and geographical distribution of the crop (Bellucci et al., 2021; Rocchetti et al., 2020; Rocchetti et al., 2022).

Seeds were collected from three different field trials, two carried out at the experimental station of the Council for Agricultural Research and Economics – Cereal and Industrial Crops (CREA-CI) in Osimo, Ancona, central Italy (latitude 43°27′ N, longitude 13°29′ N) in 2020 and 2021 and the remaining one at the ICARDA Terbol station located in Lebanon’s Bekaa Valley (latitude 33°49′ N, longitude 35°59′ E) in 2022. Table S8 report the list of lines used in the present study along with passport information and seed sample used for extraction of metabolites.

Field trials carried out in Italy were based on a Randomized Complete Block Design (RCBD) with three replicates. Each plot was constituted by a single row of 10 plants with 10 cm of distance among plants. Sowing was carried out during the first decade of April (9th-10th April), resulting in a spring-summer crop growing cycle; 480 459 chickpea lines were grown in 2020 and 2021 field trials, respectively. The field trial carried out in Lebanon was based on RCBD with two replicates. Each plot consisted of two rows of seven plants each with 25 cm of distance between plants and 50 between rows. The sowing date was 30th December 2021. Overall, 509 chickpea lines were grown. Thermopluviometric graphs of the three field trials are reported in Figure S7.

### Plant Material of the full life-cycle experiment

20 chickpea accessions were selected based on different identified QTL associated with metabolic traits (Table S5). Chickpea seeds were surface-sterilized with 70% ETOH for 3 min and then washed thoroughly with sterile water 2 times. The seeds were placed in glass petri dishes with sterile wet filter paper (soaked in sterile deionized water) for germination. The plates were kept at 22°C in the dark for germination. The seedlings with a radicle length of approximately 30 mm were used and moved in plastic pot with soil. Each plant was grown total randomized in 6 replicates in the polytunnel facilities of the MPI-MP, Golm. Each plant was fertilized twice in total, the first time when they are transferred to the pot and the second time after flowering. Plant tissue samples were collected from three periods during the experiment. The first is the vegetative stage (V), leaf, stem and root samples were collected after 4 weeks. The Second is Flowering stage (F), leaf, stem, root, flower and younger pot samples were collected after 7 weeks. The last is Senescence stage (S), leaf, stem, root and matured pot samples were collected after 10 weeks. Harvest part of the seeds after the plants were fully mature for backup and quality control.

### Metabolite extraction

Polar metabolites were extracted as described in (Salem et al., 2020). Briefly, 1 ml of pre-cooled (−20°C) methyl tert-butyl ether: methanol (3:1 v/v) was added to homogenized tissues. After, tubes were thoroughly vortexed for 1 min and then incubated on an orbital shaker (1000 rpm) for 15 min at 4°C followed by sonication for 15 min. For phase separation, a volume of 500 μl of water: methanol (3:1 v/v) was added to each tube and the samples were again thoroughly vortexed for 1 min. After that, the samples were centrifuged at 13300 g for 5 min. Semi polar phase is used for specialized metabolite analysis. Dried aliquots were resuspended in water: methanol (1:1) for specialized metabolites. Semi-polar metabolite analysis was performed on a Thermo Q Exactive Focus (Thermo Fisher Scientific, Waltham, MA, USA) with a reverse phase C18 column held at 40°C with a flow rate of 400 μl min−1 with gradual changes of eluent A (water) and eluent B (acetonitrile), both stabilized with 0.1% formic acid. Internal standards (isovitexin) were added in the extraction buffer to correct for technical variation in analytical platform.

### Chromatogram processing & compound annotation

Chromatogram processing includes chromatogram alignment based on retention time shifts, peak detection, noise filtering and isotope clustering using MS Refiner (Expressionist 12.0), samples were normalized to sample weight and batch-wise using the QC samples. Metabolite annotation was performed on pool samples analyzed in positive and negative mode with further data dependent MS2 (ddMS2) fragmenting of the top three most abundant *m/z* features for each scan. Common neutral losses were considered in reconstructing the molecular structure of the specialized metabolites, and previously reported annotated metabolites in legumes (Bulut et al., 2023).

### Genome-wide Association mapping

Genomic data were the same used by Rocchetti et al. (2024) with slightly different filtering parameters applied to the initial SNP file characterized by 6,406,472 variants in total. Briefly, variants were filtered specifically for GWA to retain only chromosomal and biallelic sites, to apply a minor allele frequency > 0.05 and to apply a maximum missing value of 0.5 by using the VCFtools (Danecek et al., 2011) commands –remove-indels –max-missing 0.9 –mac 3 –minDP 3 –maf 0.05. This narrowed the list to 701,899 SNPs). Correction for population structure was performed using the Centered Identity by state (IBS) method (Endelman and Jannink, 2012) implemented in Tassel 5. For the Association mapping mixed linear model (MLM) (Zhang et al., 2010) was used in “rMVP” (Yin et al., 2021) and linkage disequilibrium was calculated using R2, which considers the allele frequency compared to D’. The statistical significance threshold is set according to the Bonferroni corrected p-value of 0.05 for multiple testing of approximately 3.6 million times (adjusted p-value = 1.38e-8). The genome wide association mapping was performed in R.

### Linkage disequilibrium Analysis

Linkage disequilibrium (LD) was estimated using standardized disequilibrium coefficients (D), and squared allele-frequency correlations (r2) for pairs of SNP loci were determined according to the TASSEL 5 program. TASSEL 5 was also used to identify SNP-trait associations by generating a general linear model.

### Phylogenetic Analysis

Phylogenetic analysis of different gene families from different species. The amino acid sequences were aligned using from Phytozome 13. The approximately maximum-likelihood tree was constructed using aligned full-length amino acid sequences (Clustal Omega). The NJ tree and the phylogenetic tree were visualized by iTOL (https://itol.embl.de/).

### RNA-seq and data analysis

Total RNA was extracted using the NucleoSpin RNA Plant Kit (MACHEREY-NAGEL), quantified with the Qubit RNA BR Assay Kit (Thermo Fisher), and integrity was assessed on 22 samples using the RNA 6000 Nano Kit (Agilent Bioanalyzer), yielding RIN values between 7.0 and 9.7. 3′-end RNA-seq libraries were prepared from 450 ng of RNA using the Mercurius BRB-seq kit (Alithea Genomics), with 10 PCR cycles. Library quality and quantity were assessed via Agilent 4150 Tape station and KAPA qPCR. Sequencing was performed on NovaSeq6000 (150PE), producing ∼670M fragments per pool (∼8.1M/sample). Reads were aligned to the chickpea NCBI RefSeq (GCF_000331145.1) and CDCFrontier v.11 genomes using STAR v2.7.10b (Dobin A et al, Bioinformatics. 2013). Aligned reads were demultiplexed (Picard), deduplicated (UMI-tools v1.0.1), and gene-level counts obtained with FeatureCount v1.6.4 (Liao et al., 2014). Raw counts were normalized to CPM using edgeR (v3.32.1).

### cDNA cloning and vector construction

Full-length coding sequences of genes or proteins were cloned from a cDNA pool generated from chickpea leaf samples by PCR-based Gateway BP cloning using the pDONR207 and pDONR221 Donor vector (Thermo Fisher Scientific, Waltham, MA). Expression vectors for BiFC were constructed using the Gateway LR reaction with pK7FWG2 (Mansour Karimi et al., 2002; Zhang et al., 2019).

### Agrobacteria transformation and infiltration

Frozen stocks of AGL1 were added to YEB medium with Carb+ (carbenicillin 25 mg/L) and Rif+ (rifampicin 20 mg/L) and incubated at 28°C overnight. The agrobacteria were centrifuged for 30 s at 11000 rpm, 4°C and washed with 1 ml and 500 ml ice-cold water. The cells were finally resuspended in 200 ml of water (hereafter referred to as agrobacteria-competent cells). Around 1–5 ml of expression plasmid was added into a 2- ml tube with 45 ml of agrobacteria-competent cells on ice for 5 min. The solution was electrically shocked in cuvettes and kept at 28°C with 1 ml YEB medium. After shaking at 250 rpm for 1–2 h at 28°C, the cells were plated on a YEB plate (Carb+, Rif+, and appropriate antibiotics) and incubated at 28°C for 2–3 days.

The agrobacteria were grown on YEB-induced medium plates at 28°C for 36h. The cells were scraped and resuspended in 500 ml washing solution (10 mM MgCl2, 100 mM acetosyringone). After briefly vortex, 100-ml resuspended agrobacteria were diluted 10 times to measure the OD600 (should be equal to or greater than 12). The agrobacteria were finally diluted to an OD600 of 0.5 in infiltration solution (1/4MS [pH = 6.0], 1% sucrose, 100 mM acetosyringone, 0.005% [v/v, 50 ml/l] Silwet L-77). The agrobacteria were infiltrated into Tobacco leaves using a 1-ml plastic syringe, kept in the light to dry the leaves (1 h), and then subsequently kept in the dark for 24 h at room temperature (Zhang et al., 2020).

### Co-subcellular Localization Analysis

Constructs were transiently expressed in Tobacco leaves by agrobacteria-infiltration as mentioned above for the protein co-localization analysis. Confocal images were taken using a Leica DM6000B/SP8 confocal laser scanning microscope.

### Hairy root transformation and infiltration

Seeds (*Phaseolus vulgaris* & *Cicer arietinum*) were sterilized with a 30 s wash in 100% ethanol, 5 min in 0.2% sodium hypochlorite and 6 washes with sterile distilled water. After sterilization, seed were grown for 72 h at 22°C on wet filter paper and petri dishes. Then, 1-2 seeds were sown per pot (7.5 cm diameter and 9 cm tall) with vermiculite and were kept in a chamber with normal conditions (culture chamber with 300 µE.m-2s-1 lighting for 16 h at 26◦C and 8 h of darkness at 20◦C and 70% relative humidity). After 3 days in these conditions, plants were inoculated with *A. rhizogenes* K599. This inoculation was done by injection, 50-100ul of the solution with K599, at the cotyledons base. This inoculation was done by injection, 50-100ul of the solution with K599, at the cotyledons base. The inoculated part was wrapped with paper soaked in the same inoculated solution and with cling film to keep the region damp. The inoculated plants were transferred to a culture chamber with a relative humidity greater than 90%. Hairy roots appeared 7-10 days later. At this moment, the initial root was cut and the plants were transplanted again but, in that case with the transformed root and were irrigated as normal conditions. Finally, samples were collected after 2 weeks. Injection solution: *A. rhizogenes* K599 with construction should be cultivated in solid LB + kanamycin 24 hours at 28°C. Then, add 2ml of sterile distilled water and collect the bacterial suspension. Nutritive solution to irrigate the plants (3 days per week) consisted of 0.5ml/L solution (Solution A: CaCl_2_ 2 M. Solution B: KH_2_PO_4_ 1 M. pH 7. Solution C: ferric citrate 120 mM keep on darkness; Solution D: MgSO_4_ 0.5 M; K_2_SO_4_ 0.5 M; MnSO_4_ 2 mM; H_3_BO_3_ 4 mM; ZnSO_4_ 1 mM; CuSO_4_ 4 mM; CoSO_4_ 0.2 mM y Na_2_MoO_4_ 0.2 mM previously sterilized) and KNO_3_ 8mM.

## Supplementary files

Table S1 List of metabolite features identified or annotated in this research.

Table S2 Complete GWAS result showing all significant SNPs associated with the metabolic traits in chickpea seeds across three experiments.

Table S3 GWAS result showing all significant SNPs associated with the metabolic traits in two chickpea seed types.

Table S4 GWAS result showing all significant SNPs associated with the phenotypic traits in chickpea.

Table S5 List of the selected accessions used in the full life-cycle experiment.

Table S6 List of the filtered metabolite features in transient tobacco leaves.

Table S7 List of the annotated soyasaponins in transient hairy roots.

Table S8 Passport information of all the chickpea lines used in this research.

Figure S1 Genome distribution of significant SNPs related to all metabolites in two chickpea seed types

Figure S2 Genome-wide detection of selected phenotypic traits in chickpea seeds.

Figure S3 Comprehensive analysis of metabolite diversity of 20 chickpea accessions in the full life-cycle experiment.

Figure S4 Dynamic variation of soyasaponin content in different chickpea tissues across the full life-cycle.

Figure S5 Metabolic changes in transient tobacco leaves.

Figure S6 Soyasaponin changes in transient common bean hairy roots.

Figure S7 Thermopluviometric information of the three field trials

## Acknowledgments

SA, ARF, EB and RP acknowledge the financial support from the EU Horizon 2020 research and innovation Programme, project INCREASE (grant agreement No. 862862). ARF and S.A. acknowledge BG16RFPR002-1.014-0003-C01 project, financed by the European Regional Development Fund through the Bulgarian Program for Research, Innovation, and Digitalisation for Smart Transformation (PRIDST) Operational Programme. SA acknowledges the NATGENCROP project: HORIZON-WIDERA-2022- TALENTS-01, No. 101087091. SY acknowledge the financial support from China Scholarship Council.

## Author Contribution

SA conceived and designed the study. SY and SA wrote the manuscript. CS, VD, TI and AH carried out field trails and provided seed materials. EB and RP provided genome sequencing and SNPs data. SY, SA, and RW preformed metabolome experiments and data analysis. SY conducted biological validation. EB,CS, ARF and RP contributed to the editing of the manuscript. All authors contributed to the revision process and approved the final version of the manuscript.

